# Inferring the relative contributions of evolutionary processes shaping X chromosome dynamics in the common marmoset (*Callithrix jacchus*) in the presence of twinning and hematopoietic chimerism

**DOI:** 10.64898/2026.08.01.742247

**Authors:** Vivak Soni, Cyril J. Versoza, Devangana Shah, Susanne P. Pfeifer, Jeffrey D. Jensen

**Author notes:** co-corresponding authors; jointly supervised the project (;).

## Abstract

The common marmoset (*Callithrix jacchus*) is a biomedically important species that is characterized by two unusual biological traits — a high frequency of twin births and hematopoietic chimerism — that preclude the application of many commonly used population genomic approaches for quantifying evolutionary processes. In this study, we directly account for both factors in order to estimate fine-scale mutation and recombination rate maps, as well as to infer the demographic and selective processes shaping variation, on the common marmoset X chromosome. Comparing our findings to estimates recently inferred on the autosomes of this species, we find reduced rates of mutation and recombination on the X, as expected. Furthermore, population sex ratios are inferred to be nearly equal, and the appropriately rescaled autosomal population history fits the X chromosome well. Finally, we report evidence of recent selective sweeps targeting a number of X-linked genes, including several of significant biomedical relevance. Overall, these analyses provide novel insights into the evolutionary processes shaping X chromosome evolution in this biomedically-relevant primate model.

## INTRODUCTION

A number of factors shape the evolutionary trajectories of sex chromosomes in ways that are distinct from the autosomes, ranging from the age of the sex chromosome system itself (Lahn and Page 1999) to the population-level sex ratio (see the reviews of Ellegren 2011; Bachtrog et al. 2014). In most primates, the X chromosome in particular is present as a single copy in males (XY) and a double copy in females (XX). As such, under equal sex ratios, X chromosomes spend two-thirds of their evolutionary history in females. This in turn contributes to reduced levels of genetic diversity on the X chromosome relative to the autosomes, given the lower mutation rate in females (e.g., Goldmann et al. 2016; Wong et al. 2016; Jónsson et al. 2017b; Bergeron et al. 2021; Versoza et al. 2025, 2026; Terbot et al. 2025) resulting from male-biased patterns in mutation. While this may be expected from first principles due to differences in germ cell divisions in males and females (Haldane 1946; and see the reviews of Hurst and Ellegren 1998; McVean 2000), recent evidence in fact suggests that many mutations do not track cell divisions (Goldmann et al. 2016; Jónsson et al. 2017a; Gao et al. 2019; Wu et al. 2020). As there are fewer copies of the X chromosome in the population relative to an autosome (75% under even sex ratios), this results in an expected reduction in effective population size (*N_e_*) on the X, which further decreases neutral variation and necessitates alternative demographic scaling relative to the autosomes (e.g., Pool and Nielsen 2007; Singh et al. 2007). Additional factors are expected to contribute to these differences; for example, demographic patterns on the X chromosome may be affected still further by sex-biased migration (Laporte and Charlesworth 2002). Patterns of selection are also affected by the unique history of sex chromosomes relative to autosomes. For example, in an XY system, mutations occurring on the X chromosomes will be directly exposed to selection in males, regardless of dominance (Haldane 1924; Charlesworth et al. 1987; and see Bachtrog et al. 2009; Veeramah et al. 2014). A further consequence of males only carrying a single copy of the X chromosome is that recombination largely occurs only in females, resulting in a reduced overall per-generation recombination rate (e.g., Kong et al. 2002) and thus potentially increased levels of linkage disequilibrium (LD) across the X chromosome. Furthermore, increased background selection (BGS) effects can result from this decrease in recombination rate (Charlesworth et al. 1993; Charlesworth 2012).

The majority of studies of primate sex chromosome evolution have focused on the great apes as well as a few biomedically-relevant species (e.g., Osada et al. 2021; Makova et al. 2024). This inference is beginning to expand with greater data availability however; for example, a recent study of aye-ayes examined sex chromosome dynamics in this strepsirrhine of conservation concern (Terbot et al. 2025; and see Soni et al. 2025a). Amongst the emerging patterns of interest from these primate X chromosome studies, a reduced deviation from the expected 0.75 ratio of autosomal to X-linked genetic diversity has been observed in humans (e.g., Keinan et al. 2009), whilst this ratio was found to be particularly low in gorillas and orangutans (Prado-Martinez et al. 2013).

In this study, we contribute to this examination of the wider primate clade, and infer evolutionary dynamics on the X chromosome of the common marmoset (*Callithrix jacchus*). A platyrrhine native to east-central Brazil (though now found across the country due to the pet trade; Rylands and Faria 1993; Rylands et al. 2009; Garber et al. 2019), the small stature and high reproductive capabilities of the common marmoset have contributed to their use as a model for biomedical research, particularly for the study of infectious disease dynamics (owing in part to reduced levels of genetic diversity in their major histocompatibility complex relative to other mammals; Antunes et al. 1998; Wu et al. 2000; Carrion and Patterson 2012) and human neurodevelopmental disorders (e.g., Miller et al. 2016; Philippens and Langermans 2021). Importantly however, the high frequency of twin-births together with hematopoietic chimerism sets *C. jacchus* apart from the majority of other primate species, violating many of the assumptions underlying common approaches for conducting population genetic inference.

Hematopoietic chimerism refers to a phenomenon by which non-germline tissues sampled from a single individual contain genetic material from both the sampled individual itself as well as from their twin sibling (Hill 1932; Wislocki 1939; Benirschke et al. 1962; Gengozian et al. 1969). Although Ross et al. (2007) identified chimerism in a wide range of tissue types in common marmosets, Sweeney et al. (2012) hypothesized that this phenomenon was driven by blood infiltration. This explanation was recently validated by del Rosario et al. (2024), who confirmed that blood samples carry a higher proportion of sibling nuclei than other examined tissue types. A critical implication of these findings is that any non-germline tissue sampled from an individual will contain a chimeric contribution from its twin sibling, meaning that selecting alternative tissue types does not circumvent the influence of chimerism on downstream genomic analyses.

Due to the species’ biomedical importance, the common marmoset genome was among the first non-human primate assemblies published a decade ago (Marmoset Genome Sequencing and Analysis Consortium 2014), and subsequent population genomic research has since focused primarily on the characterization of standing levels of genetic variation and divergence (Faulkes et al. 2003; Harris et al. 2014; Malukiewicz et al. 2014; Yang et al. 2021; Harris et al. 2023; Yang et al. 2023; Mao et al. 2024). However, only recently have the effects of chimerism and twin-births on observed variation, and on evolutionary inference, been investigated. Based upon high-coverage, whole-genome sequencing data from 15 individuals, Soni et al. (2025b) inferred a recent population history for the common marmoset incorporating these biological factors, additionally demonstrating the potential for serious mis-inference if neglected. Further work based on this dataset additionally characterized the distribution of fitness effects (DFE) of new exonic mutations, genome-wide patterns of recent positive and balancing selection (Soni et al. 2025c), and fine-scale recombination and mutation rate heterogeneity across the *C. jacchus* genome (Soni et al. 2025d), whilst evaluating the effects of chimerism and twin-births on the inference of these various evolutionary processes. Notably, all of the above analyses have been focused exclusively upon autosomal data, leaving a deeper understanding of the evolutionary processes acting on the sex chromosomes yet to be elucidated.

With regards to expected sex chromosome dynamics, previous studies have described even sex ratios in captive common marmosets (Rothe et al. 1992), and the species exhibits little sexual dimorphism (Stevenson and Rylands 1988). However, patterns of fine-scale mutation and recombination on the X chromosome, as well as chromosome-specific demographic and selective dynamics, have not been investigated. In this study, we perform inference of these population genetic processes from high-coverage sequencing data of X chromosomes sampled from 10 unrelated individuals, and contrast these results with recent autosomal findings in the species.

## RESULTS AND DISCUSSION

### Population genomic data

Taking advantage of recent population genomic data from 15 unrelated common marmosets (7 females and 8 males; Soni et al. 2025b), we mapped the high-coverage reads of each individual to the common marmoset genome (GenBank: GCA_011100555.2; Yang et al. 2021). Based on sex-specific differences in read coverage observed in these mappings, we identified the pseudoautosomal boundary (Supplementary Figures S1 and S2), and subsequently masked the pseudoautosomal region (PAR) on the Y chromosome following best practices in humans (1000 Genomes Project Consortium 2015) to improve read mapping. We re-mapped the reads against this modified genome assembly, observing sequence coverage ratios in further support for the relatively even mixing of twin DNA in blood tissue (Table 1). Specifically, in the absence of chimerism, the proportion of sex-linked reads that are X-linked should be close to 0.5 in males (as males carry one X and one Y chromosome), and close to 1.0 in females (as females carry two X chromosomes and no Y chromosome). As chimerism will affect this ratio if the sex of the twin is different from that of the target individual, we chose the five males with a same-sex twin — all of whom exhibit an X-Y coverage ratio between 0.47 and 0.55 — for downstream analysis; additionally, we retained the three females with a same-sex twin — exhibiting a X-Y coverage ratio of >0.90 on average — as well as two additional females (Cjac_10 and Cjac_11) for whom the sex of the twin was unknown from the pedigree record, though the X-Y coverage ratio of 0.94 and 0.95, respectively, implies a female twin. We called variants in the X chromosome non-PAR of these 10 individuals in a sex-aware manner; as expected, levels of genetic variation were reduced on the X chromosome relative to the autosomes, with mean Watterson’s *θ*_w_ (Watterson 1975) across the X chromosome found to be 0.00038 per site (0.00013 for exonic regions), relative to 0.001 per site (0.00055 for exonic regions) observed on the autosomes (Soni et al. 2025b) (Table 2).

**Table 1.**
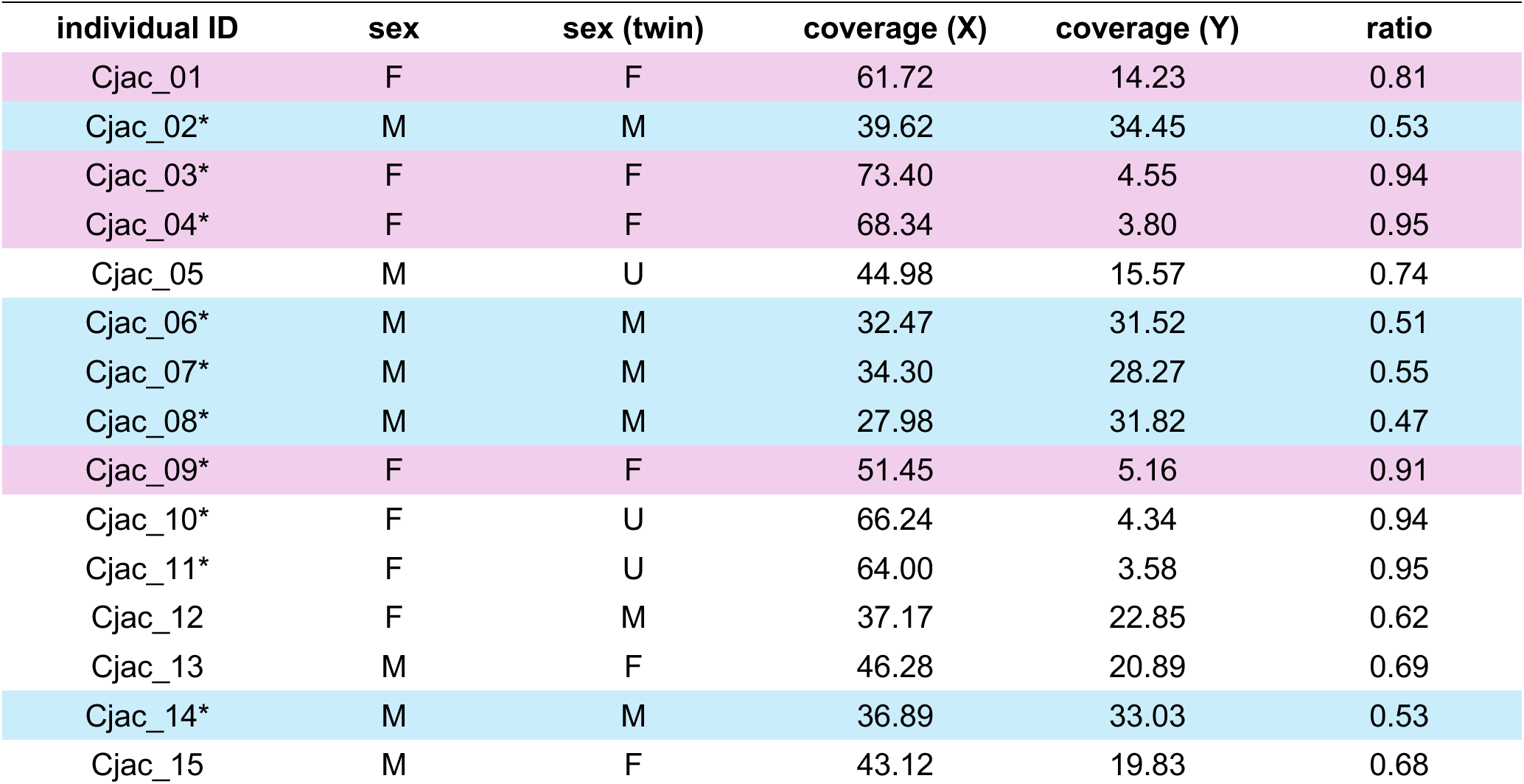
XY sequence coverage ratios of the sampled individuals. Coverage of the X and Y chromosomes as well as the XY coverage ratio is shown for the 15 unrelated common marmoset individuals sequenced by Soni et al. 2025b. The sex of the sampled individual (sex) and its twin (sex twin) is given according to the pedigree record (F = female, M = male, and U = unknown sex). Rows color-coded in pink and blue indicate females and males with same-sex twins, respectively. Individual IDs marked with a * indicate the individuals included in this study.

**Table 2.** Comparison of Watterson’s *θ_w_* and Tajima’s *D* across neutral regions of the X chromosome and the autosomes. Summary statistics were calculated across 10kb windows.

| | $\theta_w$ | | Tajima's $D$ | |
| --- | --- | --- | --- | --- |
|  | X | autosomes | X | autosomes |
| **mean** | 3.80E-04 | 1.00E-03 | -0.06 | 0.26 |
| **std** | 5.08E-04 | 9.36E-04 | 1.04 | 1.15 |
| **min** | 0 | 2.98E-04 | -2.33 | -2.72 |
| **max** | 1.04E-02 | 2.70E-02 | 2.74 | 3.77 |

### Inference of fine-scale mutation rates along the X chromosome

We combined the high-quality marmoset X chromosome generated as part of the G10K Vertebrate Genomes Project (Yang et al. 2021) used in this study with 238 other primate genomes included in the multi-species alignment of Kuderna et al. 2024. We then calculated divergence by extracting substitutions along the marmoset lineage (i.e., between *C. jacchus* and the ancestral PrimateAnc232 genome). We found that mean neutral divergence — where neutral sites were identified as those farther than 10 kb from functional regions of the genome — on the X chromosome was reduced relative to the autosomes (0.95 × 10^−3^ vs 1.31 × 10^−3^ per site, respectively, with divergence per site calculated across 100kb windows, with a minimum accessible length threshold of 1kb). We also estimated exonic divergence, finding that it was reduced relative to the maximum observed neutral divergence (see Supplementary Figure S3), as is to be expected due to the pervasive effects of purifying selection.

To calculate mutation rates, we divided the neutral X chromosome divergence by the estimated per-generation divergence time between *C. jacchus* and its nearest neighbor, *C. kuhlii*, utilizing a range of possible divergence time (0.59 million years ago [mya], 0.82 mya, and 1.09 mya; Malukiewicz et al. 2021) and generation time estimates (1.5 years and 2.0 years; Gage 1998; Tardif et al. 2003; Okano et al. 2012; Schultz-Darken et al. 2016; Han et al. 2022). Across window sizes of length 1kb, 10kb, 100kb and 1Mb, and a range of accessibility thresholds (i.e., the minimum number of accessible sites that a window must contain for it to be considered; see “Materials and Methods” for details), we inferred mean neutral X chromosome mutation rates between 0.93 × 10^−9^ per base pair per generation (/bp/gen) at the fine (1kb) scale (assuming a divergence time between *C. jacchus* and *C. kuhlii* of 1.09 mya and a generation time of 1.5 years) and 4.05 × 10^−9^ /bp/gen at the large (1Mb) scale (assuming a divergence time between *C. jacchus* and *C. kuhlii* of 0.59 mya and a generation time of 2.0 years) (see Figure 1a for density plots of mutation rates across generation and divergence times and Supplementary Table S1 for the full range of mean rates). As evident from the fine-scale mutation rate map shown in Figure 1b, the sex-averaged mean mutation rate across the X chromosome for the 2.0-year generation time (2.3 10^−9^ /bp/gen) is reduced relative to the autosomal rate (3.2 × 10^−9^ /bp/gen); this rate reduction relative to the autosomal fine-scale mutation rate is consistent with the reduction observed in humans, where the mutation rate on the X chromosome is 70–75% of that inferred on the autosomes (Nachman and Crowell 2000; Makova and Li 2002; and see the review of Schaffner 2004). Notably, these inferred rates — both on the common marmoset X chromosome and the autosomes — are somewhat lower than the direct pedigree-based mutation rate estimate of 4.3 10^−9^ /bp/gen inferred by Yang et al. (2021), though chimerism was unaccounted for in that study (chimerism may be expected to bias pedigree-based estimates of mutation rates given that chimeric samples contain DNA from the sampled individual as well as its twin, which may in turn affect counts of *de novo* mutations observed in parent-offspring trios). Our inferred rate is also lower than estimates in other primates (e.g., Bergeron et al. 2023), and less than half of that inferred in coppery titi monkeys (Soni, Versoza et al. 2026; Versoza et al. 2026) and owl monkeys (Thomas et al. 2018), two fellow platyrrhines. Finally, following from Miyata et al. (1987), one may consider male/female ratios of mutation rates as (4-3*R*)/(3*R*-2), where *R* is the X to autosome ratio of mutation rates noted above, which in turn implies a roughly 10-fold male bias.

**Figure 1.**
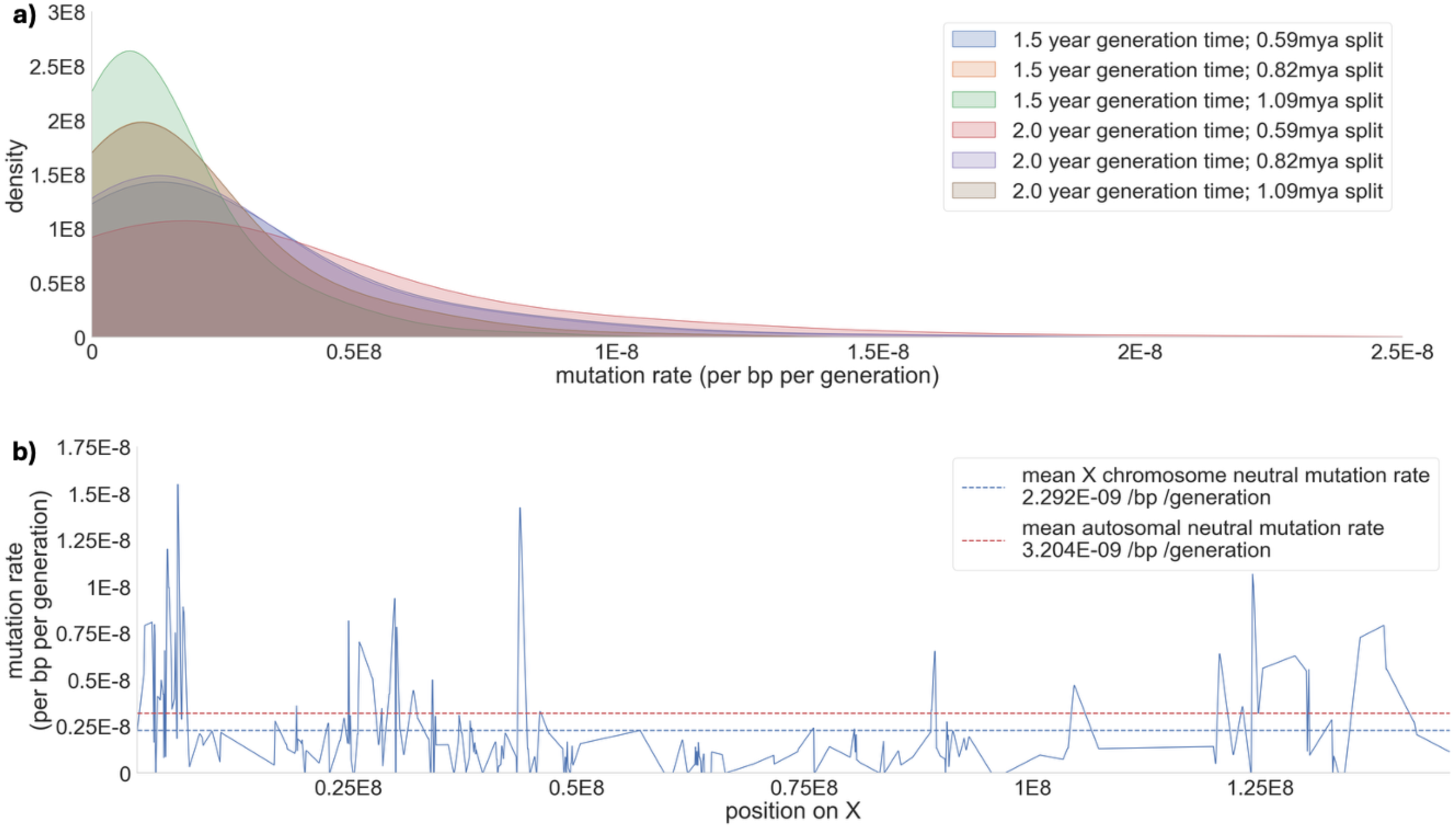
Fine-scale mutation rates along the X chromosome. **a)** Density plots of the per-site per-generation mutation rate implied by the neutral divergence between *C. jacchus* and the closely related *C. kuhlii* for three divergence times (0.59 mya, 0.82 mya, and 1.09 mya; Malukiewicz et al. 2021) and two generation times (1.5 years and 2.0 years; Gage 1998; Tardif et al. 2003; Okano et al. 2012; Schultz-Darken et al. 2016; Han et al. 2022), across 100kb windows with an accessibility threshold of 1kb. **b)** Landscape of mutation across the X chromosome for windows of size 100kb, assuming a divergence time between *C. jacchus* and *C. kuhlii* of 0.82 mya and a generation time of 2.0 years, across 100kb windows with an accessibility threshold of 1kb. The blue dashed line indicates the mean neutral mutation rate across the X chromosome whereas the red dashed line indicates the mean neutral mutation rate across the autosomes using the same window sizes and accessibility thresholds, as inferred by Soni et al. (2025d).

### Inference of the population history of the X chromosome

To characterize population structure on the common marmoset X chromosome, we ran ADMIXTURE (Alexander et al. 2009) for values of *k* (the number of demes) from 1 to 5, with a single population identified as the most likely (*k* =1, cross validation error [CVE] = 1.35; *k* =2, CVE = 2.12; *k* =3, CVE = 2.43; *k* =4, CVE = 3.06; *k* =5, CVE = 3.15). This result is consistent with the findings of Soni et al. (2025b) pertaining to the analysis of population structure on the autosomes, as well as with the localized geographic range of *C. jacchus*, characterized by little fragmentation and bounded by major river systems (see Figure 3 of Malukiewicz et al. 2020).

Given this consistency with the autosomes in terms of population structure, we evaluated the fit of the common marmoset autosomal demographic history (Soni et al. 2025b) — which models both twin-birthing and hematopoietic chimerism — to the X chromosome under a range of sex ratio scalings. Specifically, we simulated 100 replicates of a 1Mb neutral region under a demographic model consisting of an ancestral population of ∼62k individuals that experienced a bottleneck ∼3,500 generations ago, reducing the population to about a third of its size, before gradually recovering to its current day size of ∼34k individuals (Soni et al. 2025b), with sex-specific mutation rates across multiple male:female sex ratios (0.3:0.7, 0.4:0.6, 0.45:0.55, 0.5:0.5, 0.55:0.45, 0.6:0.4, 0.7:0.3), and compared the fit of Watterson’s *θ*_w_ (Watterson 1975) as a measure of genetic variation and Tajima’s *D* (Tajima 1989) as a summary of the site frequency spectrum with neutrally evolving regions in the empirical data (see “Materials and Methods” for further details). As shown in Figure 2, male:female sex ratios ranging from 0.4 to 0.6 best fit the empirical data, with the alternative sex ratios generally reducing variation below the empirical observation. Notably, expected variation decreases as the sex ratio deviates further from 0.5:0.5 in either direction. This is explained by the fact that males have a higher mutation rate than females, but fewer X chromosomes. Thus, as the male sex ratio increases, so does the mean mutation rate per site, whilst the *N_e_* of the X decreases. Conversely, at higher female sex ratios, the mutation rate per site is reduced whilst the *N_e_* of the chromosome is generally increased (Supplementary Table S2). These relative differences between sex ratios are consistent with simulated expectations (see Figure 1 of Spurley and Payseur 2025). Furthermore, the *N_e_* on the X chromosome is expected to be 25% lower than on the autosomes under even sex ratios owing to chromosome count alone. Given that the estimated autosomal *N_e_* (calculated from Watterson’s *θ*_w_ inferred by Soni et al. [2025b] and the fine-scale mutation rate inferred by Soni et al. [2025d]) is 78,027, the expected *Ne* on the X chromosome would be 58,520 if this were the only contributing factor. Supplementary Table 2 compares these expected *N_e_* reductions based on empirical *θ*_w_, and serves to further support a roughly equal sex ratio (as previously observed in common marmoset colonies; Rothe et al. 1992).

**Figure 2.**
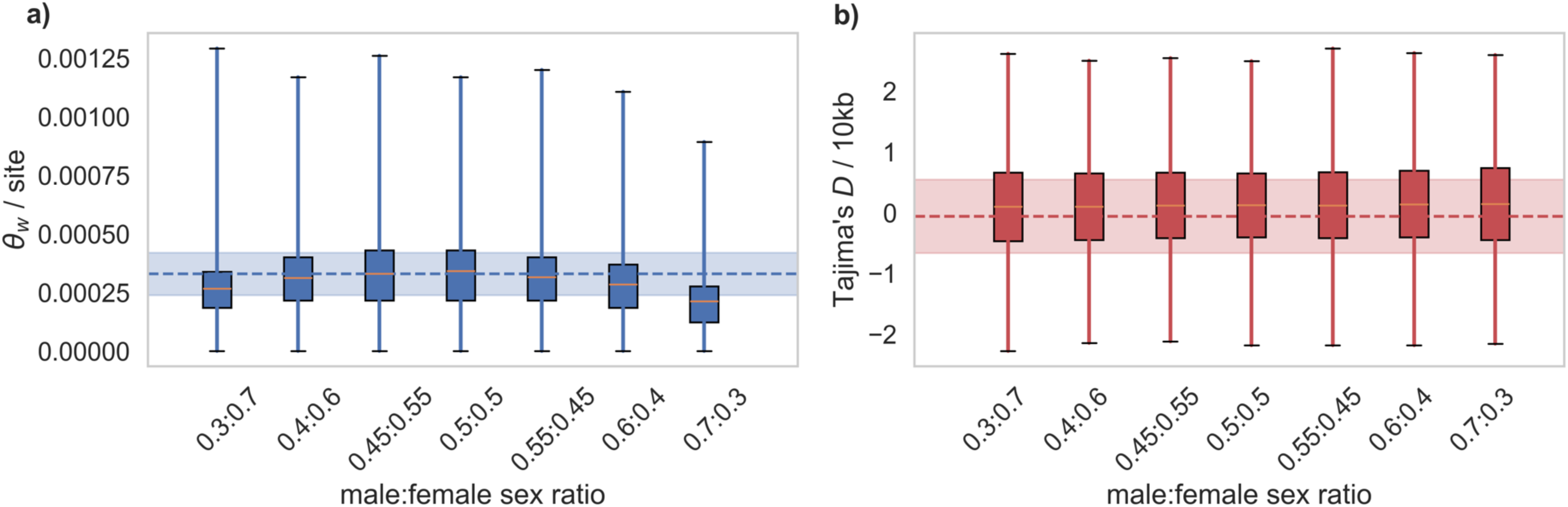
Population genetic summary statistics from the inferred population history of the X chromosome for varying male:female sex ratios. **a)** Fit of Watterson’s *θ*_w_ (shown in blue; Watterson 1975) and **b)** Tajima’s *D* (shown in red; Tajima 1989) between the mean values observed in the empirical data (dashed lines) and the simulated data (box plots). Shaded regions indicate the median absolute deviation with a normal distribution scaling factor. Box plots indicate the median (orange line), the 25^th^ and 75^th^ percentiles (boxes), and the minimum and maximum values (black lines) across 100 simulation replicates under the demographic model inferred by Soni et al. (2025b), rescaled to the X chromosome (see “Materials and Methods” for further details).

**Figure 3.**
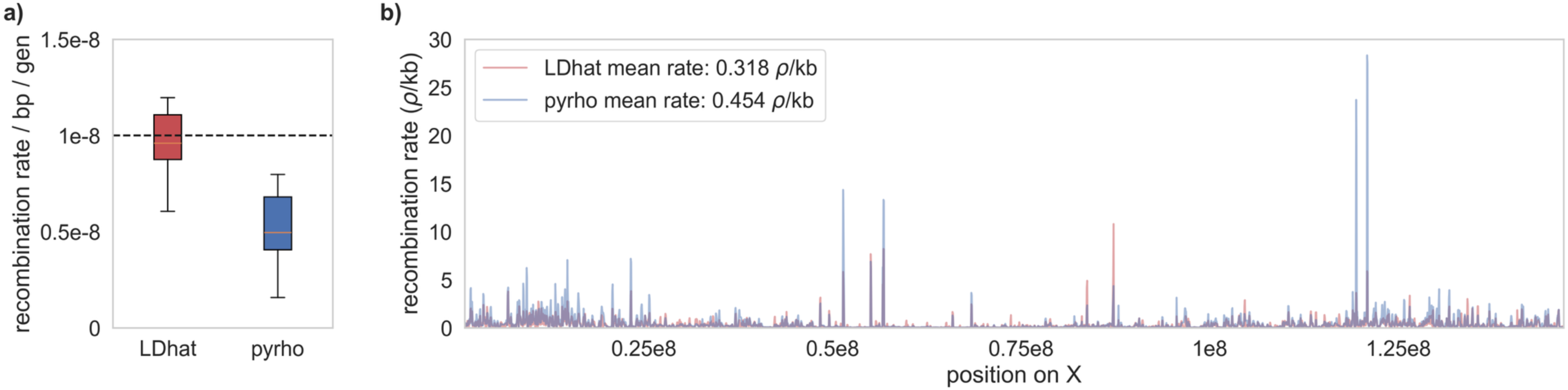
Fine-scale recombination rates along the X chromosome. **a)** Recombination rates inferred using LDhat (shown in red) and pyrho (shown in blue) from benchmarking simulations, in which 10 replicates of a 1Mb genomic region were simulated under the demographic model inferred in this study, assuming a fixed recombination rate of 1 × 10^−8^ /bp/gen (dashed black line). Box plots indicate the median (orange line), the 25^th^ and 75^th^ percentiles (boxes), and the minimum and maximum values (solid black lines). **b)** Fine-scale recombination maps inferred using LDhat (shown in red) and pyrho (shown in blue) across 100kb windows along the X chromosome, for mutation rates calculated using a generation time of 2.0 years with rates rescaled by the extent of mis-inference observed from benchmarking analysis (panel **a**).

Finally, Table 2 compares these summary statistics across neutral regions of the X chromosome and the autosomes. As shown, there is a strong reduction in the levels of neutral variation on the X chromosome (∼62% lower) — a reduction considerably more severe than the 25% expected by chromosome count alone. Thus, the reduced mutation rate on the X, combined with greater expected reductions in variation owing to purifying selection in this reduced recombination environment as described below (also supported by a stronger skew towards rare alleles on the X), are all likely additional contributing factors.

### Inference of fine-scale recombination rates along the X chromosome

We used both the demography-unaware estimator LDhat (McVean et al. 2002, 2004; Auton and McVean 2007) and the demography-aware estimator pyrho (Spence and Song 2019) to infer fine-scale recombination rate maps on the common marmoset X chromosome. Prior to performing inference on the empirical data, we evaluated how both estimators perform under the inferred *C. jacchus* X chromosome demographic model (see “Inference of the population history of the X chromosome”) — an evaluation that is particularly important in this species, given that frequent twin-birthing and chimerism violate assumptions of the underlying model, and may be expected to themselves generate LD. To this end, we simulated 10 replicates of a 1Mb neutrally evolving genomic region under the common marmoset demographic model including twinning and chimerism, with a fixed recombination rate of 10^−8^ /bp/gen, and then inferred recombination rates using LDhat and pyrho to evaluate the extent of mis-inference. As shown in Figure 3a, both methods underestimated the recombination rate under this scenario. These results are consistent with those of a recent simulation study of Dutheil (2024), which found that population contractions and expansions (both of which have been inferred in the recent history of the common marmoset population) often result in an underestimation of the recombination rate when performing inference using LD-based approaches. Furthermore, Soni et al. (2025d) found that LDhat estimates a higher recombination rate relative to pyrho under twin-birthing and chimerism — a trend which was borne out in our benchmarking analysis as well.

With these benchmarking results in hand, we performed inference with both estimators on the empirical data, and re-scaled the results based on the extent of expected mis-inference as quantified in the simulation analysis. Encouragingly, the resulting mean population-scaled recombination rates inferred by LDhat and pyrho are in reasonably close agreement (0.318 ρ/kb and 0.454 ρ/kb; Figure 3b). These rates observed on the X chromosome are substantially lower than the mean inferred autosomal rate of 0.895 ρ/kb (Soni et al. 2025d), giving a ρ*_x_*/ρ*_a_* ratio (i.e., the ratio of the X chromosomal to autosomal recombination rate) of 0.36 and 0.51 respectively. For comparison, summarizing in cM/Mb, Kong et al. (2002) reported a ratio of 0.81 in humans.

### Scans for recent positive selection and balancing selection on the X chromosome

It has previously been shown that outlier approaches (i.e., approaches that identify candidate loci from an arbitrarily chosen tail of a genomic distribution — often 5% or 1%) for performing genomic scans for selection are frequently associated with high false-positive rates (Teshima et al. 2006; Thornton and Jensen 2007; Jensen et al. 2008; Crisci et al. 2012; Jensen 2023; Soni et al. 2023). We therefore constructed an evolutionarily appropriate baseline model to account for the species-specific mutation and recombination environment, demographic history, purifying and BGS effects in functional regions, and reproductive dynamics, as recommended by Johri et al. (2022a,b), in order to determine if particular loci are poorly fit by the expectations arising from these constantly-operating evolutionary processes alone. To do so, we simulated the full non-PAR of the *C. jacchus* X chromosome, modelling the empirical exonic structure, and drawing mutations in exonic regions from the DFE inferred in common marmosets by Soni et al. (2025c) comprised of four fixed classes (Johri et al. 2020): effectively neutral mutations (2*N*_ancestral_ *s* < 1, where *N_ancestral_* is the ancestral population size and *s* is the reduction in fitness of the mutant homozygote relative to wild-type), *f_0_* = 0.3; weakly deleterious mutations (1 ≤ 2*N*_ancestral_ *s* < 10), *f_1_* = 0.1; moderately deleterious mutations (10 ≤ 2*N*_ancestral_ *s* < 100), *f_2_* = 0.1; and strongly deleterious mutations (100 ≤ 2*N*_ancestral_ *s*), *f_3_* = 0.5 (see “Materials and Methods” for details). We simulated 100 replicates under this baseline model, and then performed genome scans for selective sweeps and balancing selection using SweepFinder2 (DeGiorgio et al. 2016) and the *B_0MAF_* method implemented in BalLeRMix (Cheng and DeGiorgio 2020) on this simulated data. The maximum composite likelihood ratio (CLR) value inferred by each method across all windows and replicates was set as the null threshold for the empirical inference of the respective form of selection, under the logic that this value represents the highest CLR that can be generated under the population-specific model in the absence of positive or balancing selection. The null thresholds for positive and balancing selection inference were found to be 67.40 for SweepFinder2 and 122.78 for *B_0MAF_*.

We next performed inference on our empirical data, using the same analyses as applied to the simulated data to generate null thresholds (see “Materials and Methods” for further details). Utilizing the thresholds from the evolutionary baseline model, we identified a total of 1,821 selective sweep candidates (5.7% of all tested loci) and 1,685 balancing selection candidates (5.3% of all tested loci; see Supplementary File S1 for genome scan results at each single nucleotide polymorphism [SNP] and the maximum CLR per candidate gene). Relative to the autosomes, these represent a high proportion of candidate loci (Soni et al. 2025c found that only 0.3% and 0.02% of all tested loci were sweep and balancing selection candidates on the autosomes, respectively). However, as depicted in Figure 4, the majority of candidate loci (indeed, all selective sweep candidate loci) are clustered in one location near the end of the X chromosome. This region is notable for a substantial increase in gene density (Figure 4e), though the recombination rate in this region does not substantially differ from the chromosome-wide average (Figure 4f). It is certainly feasible that a single gene, or a small number of genes, is/are being targeted by positive or balancing selection in this region, and the resulting signatures of selection are detectable across a wide genomic window. We therefore focused our analysis on genes under the peaks of the likelihood surface in this region (see Figures 4b and d).

**Figure 4:**
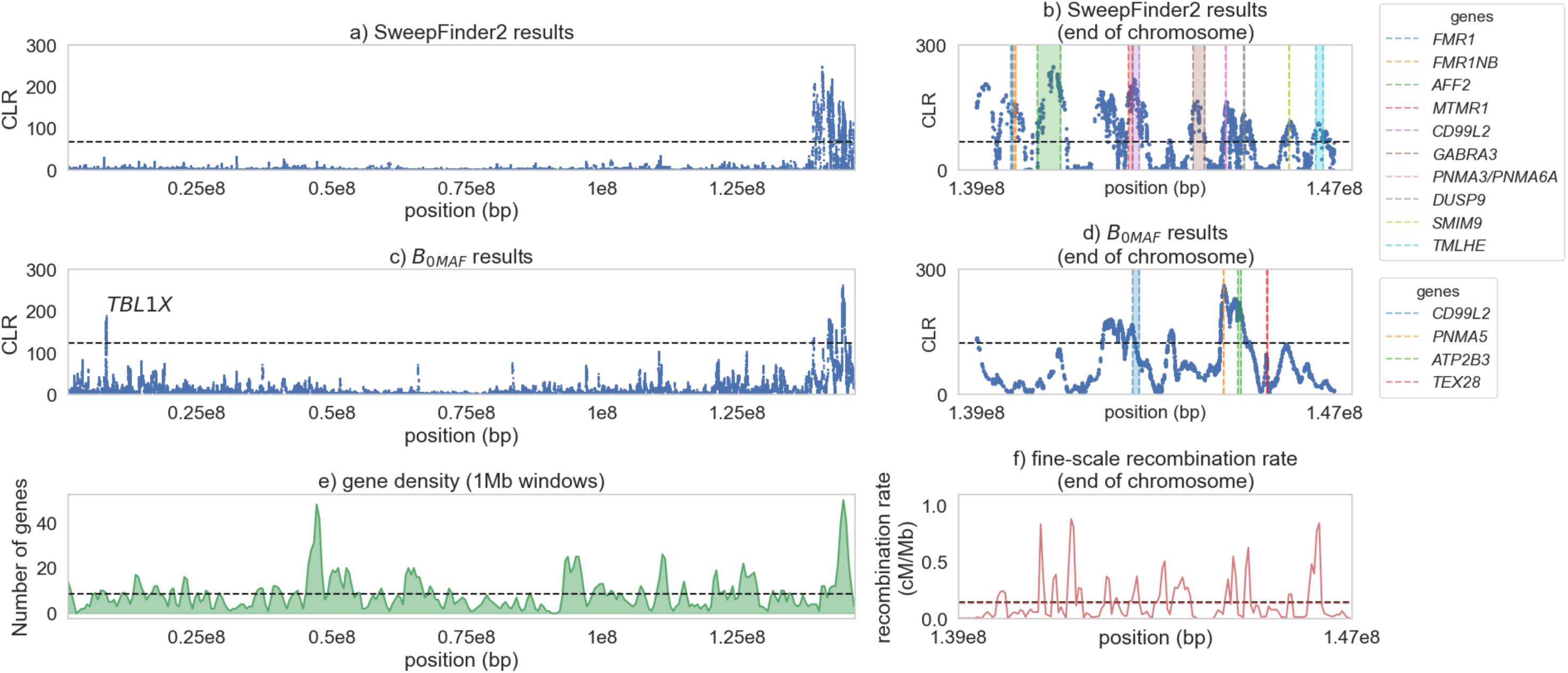
Results of genome scans for selective sweeps and balancing selection. Scans for **a)** selective sweeps performed using SweepFinder2, and **c)** balancing selection performed using *B0MAF*. In each case, the x-axis shows the position along the X chromosome, the y-axis shows the composite likelihood ratio (CLR) value of the statistic at each SNP. The horizontal dashed line represents the null threshold for detection. Panels **b)** and **d)** zoom in on the genome scan results from panels **a)** and **c)** respectively, showing from position 1.39e8 onwards, where a large number of candidate loci clustered together. For panel **b)** and **d)**, loci that map to candidate genes at the peaks of the likelihood surface are highlighted. **e)** Gene density across the common marmoset X chromosome, calculated across 1Mb windows. The dashed line represents the mean number of genes per window (8.6). **f)** Fine-scale recombination rate on the X chromosome, from position 1.39e8 onwards. The black dashed line represents the mean chromosome-wide recombination rate, whilst the red dashed (nearly overlapping) line represents the mean recombination rate across the region of the chromosome plotted in this panel.

Notably, a number of genes identified in these scans are functionally associated with neurodevelopment and neurodevelopmental disorders:

- *FMR1*, *FMR1NB* and *GARBRA3*: Both *FMR1* and its neighbor, *FMR1NB*, were identified as selective sweep candidates. The silencing of the *FMR1* gene causes Fragile X syndrome (FXS), a monogenic cause of autism spectrum disorder (ASD) and intellectual disability (Chaste et al. 2012). Harbers et al. (2025) found that female common marmosets heterozygous for an *FMR1* mutation exhibit phenotypes associated with FXS in humans, marking this marmoset model’s potential for evaluating therapeutic targets of FXS. *FMR1* has also been identified as a candidate for recent positive selection in humans by Crespi et al. (2010), who hypothesized that the cause of positive selection may be due to alterations of common neurogenetic pathways. Harbers et al. (2025) additionally found that marmoset models for FXS (i.e., those carrying the *FMR1* mutant) exhibit dysregulation of synapse-related genes. Indeed, another of our selective sweep candidate genes, *GABRA3*, was identified as one such gene exhibiting consistent changes in expression.
- *AFF2*: Candidate loci in this gene (which was previously named *FMR2*) exhibited the highest likelihood peak in our scans for selective sweeps (CLR = 248). It has previously been shown that the silencing of *AFF2* causes fragile X E (FRAXE) syndrome, a milder form of ASD compared with FXS (Gu et al. 1996). Furthermore, Mondal et al. (2012) found an excess of missense mutations at highly conserved evolutionary sites in *AFF2* in male patients with ASD. Given that ASD affects four times as many males as females, their study supports the hypothesis that recessive variants in this X-linked gene are associated with increased risk of ASD in males. Further, Zou et al. (2022) identified hemizygous missense variants in *AFF2* in males with partial epilepsy, concluding that this gene is potentially causative of X-linked partial epilepsy with antecedent febrile seizures. This finding is particularly relevant given that marmosets are a model for generalized epilepsy (Yang et al. 2022), and *AFF2* has been identified as a candidate for positive selection in human populations by both Sabeti et al. (2007) and Lambert et al. (2010).
- *TMLHE*: A hemizygote pathogenic variant in this selective sweep candidate gene has been shown to result in X-linked autism type 6, a condition characterized by moderate intellectual disability (Verhoeven et al. 2025).
- *TEX28*: This balancing selection candidate gene is frequently found in a cluster of genes that are duplicated in what is known as Xq28 duplication syndrome. This duplication is characterized by developmental delays and an increased risk of ASD (Kopytova et al. 2025).
- *PNMA3/PNMA6A* (selective sweep and balancing selection candidates), and *PNMA5/PNMA6F* (balancing selection candidates): Mutations in the *PNMA* family of genes are associated with human intellectual disability as well as developmental abnormalities (Schüller et al. 2005), and *PNMA3* has previously been described as being targeted by positive selection in primates (Cagliani et al. 2024).

Though not related to neurodevelopment, another gene with statistical support for being targeted by balancing selection was of particular interest for the biology of common marmosets:

- *ATP2B3*: The enzyme encoded by *ATP2B3* removes bivalent calcium ions from eukaryotic cells, and is crucial for maintaining intracellular calcium homeostasis, particularly in neural, brain, and adrenal tissues (Wang et al. 1994). Given that marmosets struggle to digest

this mineral and have a lack of readily available calcium in their diet (Jarcho et al. 2013), as well as the enrichment in candidate genes related to calcium having been found on the common marmoset autosomes (Soni et al. 2025c), our identification of *ATP2B3* as a candidate gene being maintained by balancing selection lends further support to hypotheses of on-going selection related to calcium intake in common marmosets.

Finally, a number of other candidate genes are associated with a range of previously described medical disorders: MTMR1 is associated with X-linked myotubular myopathy (Laporte et al. 2001), *CD99L2* is highly expressed in the brain and plays a key role in synaptic plasticity and spatial memory (Kang et al. 2025), *DUSP9* has been shown to be critical for placental development in mice (Czikk et al. 2013), *SMIM9* has been associated with congenital abnormalities such as cloacal exstrophy (Nordenskjöld et al. 2023), and *TBL1X* has been linked to pancreatic cancer and lymphomas (Pray et al. 2023).

## CONCLUDING THOUGHTS

This study provides the first insights into the evolutionary forces shaping X chromosome evolutionary dynamics in common marmosets, whilst accounting for the effects of chimerism and twin-birthing. Results suggest that both fine-scale mutation and recombination rates are reduced on the X chromosome relative to the autosomes. While the directionality of both patterns is to be expected and is consistent with findings in humans (e.g., Kong et al. 2002), the magnitude of the reduction in recombination is considerably greater in common marmosets. In addition, the rescaled autosomal population history inferred by Soni et al. (2025b) fit the X chromosome well, with simulations lending further support for roughly equal sex ratios in this population. Finally, chromosome-wide scans identified a number of genes with strong statistical support for having experienced recent selective sweeps or balancing selection. A number of these genes are related to neurodevelopment and are thus highly relevant to the role of the common marmoset as a model for biomedical research.

One of the most notable observations of this study was the level of reduction of variation on the X relative to the autosomes, with an *X/A* ratio of 0.38 (nearly half that reported in humans; see Figure 1 of Gottipati et al. 2012). However, our results provide a number of insights into the evolutionary processes driving this reduction. Firstly, as our results suggest that the autosomal demographic history with equal or nearly equal sex ratios well fits observed patterns of variation on the X, the difference in population chromosome count would be expected to reduce *N_e_* on the X by ∼25%. Secondly, the estimated mutation rate on the X was ∼71% of that observed on the autosomes, further reducing expected levels of variation. Thirdly, the mean inferred recombination rate was approximately half of that inferred on the autosomes, increasing expected diversity-reducing BGS effects across the chromosome (with exonic ratios of variation further suggesting stronger purifying selection acting on the X). Finally, unlike the majority of other primates characterized to date, twinning in marmosets results in a yet further reduction of *N_e_*, and therefore an additional reduction in population-level recombination leading to longer-range BGS effects. Thus, these factors all likely contribute to the unusual *X/A* ratio observed in this population.

## METHODS AND MATERIALS

### Population genomic data

This study was based on population genomic data of 15 unrelated common marmosets (7 females and 8 males) previously generated by Soni et al. 2025b. Following best practices in the field (Pfeifer 2017), we used SOAPnuke v.1.5.6 (Chen et al. 2018) to remove adapters (“*--cutAdaptor*”) as well as trim homopolymers (“*-A* 0.25 *-G*”), polyX tails (“*--polyX*”), and low-quality bases (“*-l* 20 *-q* 0.3 *-Q* 2”), thereby discarding any reads that contained more than 1% ambiguous bases (“*-n* 0.01”) or that were shorter than 150 bp (“*--minLen* 150”). Using BWA-MEM v.0.7.17 (Li 2013), we mapped the quality-controlled reads against the common marmoset genome (GenBank: GCA_011100555.2; Yang et al. 2021), marking any secondary hits (“*-M*”). After mapping, we marked read duplicates in each sample using the Genome Analysis Toolkit (GATK; van der Auwera and O’Connor 2020) *MarkDuplicates* v.4.2.6.1. Based on sex-specific differences in read coverage observed in these de-duplicated mappings, we identified the pseudoautosomal boundary. We then masked the PAR on the Y chromosome following best practices in humans (1000 Genomes Project Consortium 2015) to improve read mapping and variant calling on the sex chromosomes. We re-mapped the quality-controlled reads against this modified genome assembly following the approach outlined above and then called sites in the non-PAR in a sex-aware manner. Specifically, we called both variant and invariant sites using GATK’s *HaplotypeCaller* (with the “*-ERC* BP_RESOLUTION flag enabled) from high-quality read mappings (“*--minimum-mapping-quality* 40”), using a sample ploidy of 1 (“*--sample-ploidy* 1”) for males with a same-sex twin as males only carry one copy of the X chromosome; as females with a same-sex twin carry two copies of the X chromosome, their sample ploidy was set to 2 (“*--sample-ploidy* 2”). With the exception of Cjac_10 and Cjac_11 for whom the X-Y coverage ratio implied a same-sex twin, we excluded individuals with different-sex twins from further analyses given the uncertainty in the extent of chimerism in each sample. As sequencing libraries were previously prepared following a PCR-free protocol, we disabled PCR error correction (“*--pcr-indel-model* NONE”) following the developers’ guidelines (van der Auwera and O’Connor 2020). We combined the calls from each individual (GATK *CombineGVCFs*) in order to jointly genotype all samples at all sites (GATK *GenotypeGVCFs*, with the “*-all-sites*” flag enabled). To facilitate comparisons with the autosomes, we applied previously established methods to construct high-confidence datasets of fully genotyped biallelic SNPs and monomorphic sites comprising putatively neutral regions for analyses of population structure and demographic history (Soni et al. 2025b), alongside chromosome-scale datasets used to estimate mutation and recombination rates (Soni et al. 2025d) as well as patterns of selection (Soni et al. 2025c).

### Inference of fine-scale mutation rates along the X chromosome

We mirrored the approach presented in the preprint of Soni et al. (2025d) to infer fine-scale rates of neutral divergence and mutation along the X chromosome. Specifically, we replaced the common marmoset X chromosome included in the 239-way multiple primate alignment (Kuderna et al. 2024) with the high-quality version generated as part of the G10K Vertebrate Genomes Project (Yang et al. 2021) used in this study. We then extracted substitutions along the marmoset lineage (i.e., between *C. jacchus* and the ancestral PrimateAnc232 genomes), removing any alignments less than 10kb in length to account for fragmentation that might affect estimates of divergence and mutation, and masking any SNPs segregating in the species. We calculated neutral divergence across windows of size 1kb, 10kb, 100kb, and 1Mb, with a step size of 500bp, 5kb, 50kb, and 500kb respectively, masking ± 10 kb of functional regions and conserved elements to account for the effects of purifying selection and BGS. The number of accessible sites in any given genomic window was calculated as the number of sites that were left unmasked. To calculate mutation rates, we divided these neutral divergence rates by the per-generation divergence time between *C. jacchus* and its nearest neighbor, *C. kuhlii*, utilizing three possible divergence times (0.59 mya, 0.82 mya, and 1.09 mya; Malukiewicz et al. 2021) and two possible generation time estimates (1.5 years and 2.0 years; Gage 1998; Tardif et al. 2003; Okano et al. 2012; Schultz-Darken et al. 2016; Han et al. 2022).

### Inference of the population history of the X chromosome

Prior to performing demographic inference on the X chromosome, we determined the extent of population structure using the software ADMIXTURE v.1.3.0 (Alexander et al. 2009), using a range of *k* values from 1 to 5, where *k* is the number of demes. We used the value of *k* with the lowest CVE for deme assignment.

To account for the biasing effects of purifying and BGS on demographic inference (Charlesworth et al. 1993; Johri et al. 2021), we based our demographic analyses on a sub-set of the empirical data in which functional regions, as well as 10kb on the flanks of each functional region (Johri et al. 2020), were masked. Given that the number of demes was the same on both the autosomes (*k* = 1; Soni et al. 2025b) and the X chromosome, we evaluated the fit of the demographic history inferred on the common marmoset autosomes by Soni et al. (2025b) to the X chromosome. Briefly, their model involved an ancestral population of 61,198 individuals collapsing to 17,931, with the size reduction occurring 3,513 generations ago. The population was inferred to gradually recover to its current day size of 33,830 individuals. We simulated the X chromosome under this population history using SLiM 5.0.1 (Haller et al. 2026), utilizing the *initializeSex(“X”)* within our SLiM script to account for X chromosome scaling. Importantly, the demographic model of Soni et al. (2025b) additionally incorporates chimerism and twin-birthing; to do so, monogamous mating pairs produce non-identical twins, and the genotypes of these twins are combined post-simulation to artificially generate a chimeric individual (see Figure 1 of Soni et al. 2025b for further details). Following this approach, we simulated a 10*N_ancestral_* generation burn-in time (where *N_ancestral_* is the initial population size of 61,198 individuals) prior to the demographic model, to allow genetic variation to reach equilibrium, and then generated a total of 10 chimeric individuals post-simulation to mirror our empirical sample size of individuals with same-sex twins. Specifically, we simulated 100 replicates of a 1Mb region, with variable mutation and recombination rates, as it has been previously shown that ignoring heterogeneity in mutation and recombination across the genome can result in the mis-inference of population history (Soni et al. 2024). To this end, we drew mutation rates from the empirical X chromosome mutation rate map inferred in this study (see “Inference of fine-scale mutation rates along the X chromosome”). As the X chromosome spends more time in females relative to males, and as previous work in primates has demonstrated a male mutation bias of ∼2–3 in multiple species (Thomas et al. 2018; Wang et al. 2020; Wu et al. 2020; Versoza et al. 2025), we accounted for this sex-specific mutation bias by drawing a mutation rate for each 1kb window and rescaling it such that the male mutation rate was ∼2.7 times greater than that of females and the mean rate across all windows was equal to the rescaled empirical mean rate (i.e., the autosomal mean rate rescaled for males and females). Notably, as the sex-specific mutation bias has not been well-quantified in common marmosets, this estimate relies on observations in other closely-related species. We drew recombination rates from a uniform distribution, with a minimum value of 0, and a maximum value of 2 × 10^−8^ /bp/gen, such that the mean rate across each simulated 1Mb region was equal to 1 × 10^−8^ /bp/gen. In order to investigate the possibility of sex-specific population dynamics, we simulated under the following male:female sex ratios: 70:30, 60:40, 55:45, 50:50, 45:55, 40:60, and 30:70.

We compared the fit of Watterson’s *_θ_*_w_ (Watterson 1975) and Tajima’s *D* (Tajima 1989) from each sex-ratio simulation set to those obtained from putatively neutral regions in the empirical data, with each summary statistic calculated across 10kb windows with a 5kb step size using pylibseq v.1.8.3 (Thornton 2003).

### Inference of fine-scale recombination rates along the X chromosome

Following the approach of Soni et al. (2025d), we inferred fine-scale recombination rates on the X chromosome using the demography-unaware estimator LDhat v.2.2 (McVean et al. 2002, 2004; Auton and McVean 2007) and the demography-aware estimator pyrho v.0.1.7 (Spence and Song 2019). As both LDhat and pyrho are coalescent-based inference approaches that rely on the Wright-Fisher framework, we evaluated the impact of twin-births and chimerism — two processes that violate the assumption of the model — on recombination rate inference on the X chromosome. To this end, we simulated 10 replicates of a 1Mb region under the demographic model inferred in this study (see “Inference of the population history of the X chromosome” for details of parameterizations of this model), with a fixed mutation rate of 1 × 10^−8^ /bp/gen and a fixed recombination rate of 1 × 10^−8^ /bp /gen, and then estimated recombination rates from these simulated replicates using both LDhat and pyrho. To account for the extent of mis-inference observed from this simulated data, we rescaled the recombination rates inferred from the empirical data using a scaling factor of 1.04 for LDhat and 2.02 for pyrho.

### Scans for recent positive selection and balancing selection on the X chromosome

To calculate exonic divergence, we utilized the same approach described above to estimate neutral divergence (see “Inference of fine-scale mutation rates on the X chromosome”), limiting the procedure to exonic regions only.

In order to generate null thresholds for the inference of selection, we simulated 100 replicates of the total non-PAR of the common marmoset X chromosome under the demographic model outlined above (see “Inference of the population history of the X chromosome”), with an equal male:female sex ratio (as determined in this study). We directly modelled the exonic structure from the empirical data. Given the large population sizes in the common marmoset demographic model and the need to simulate forward-in-time to model chimerism and twin-birthing, simulating the entire non-PAR of the X chromosome was not tractable. As such, we split the X chromosome into segments wherever a 50kb gap existed between exons. When such a gap was identified, we set a breakpoint such that each segment ended with half of the total distance between these exons, leaving a minimum 25kb non-functional region at the end of each segment (see Supplementary Figure S4 for a schematic example). This approach ensured that we had sufficient distance from functional regions that the effects of BGS were not significant. In total, we simulated 160 segments per replicate. Given that mutation and recombination rate variation can affect the power of genome scans (Soni et al. 2023, 2024; Soni and Jensen 2024), we drew mutation rates from the empirical X chromosome fine-scale mutation rate distribution, and recombination rates from a uniform distribution, such that the mean mutation rate across each simulation replicate was equal to the mean X chromosome mutation rate, and the mean recombination rate across each simulation replicate was equal to the mean fine-scale recombination rate inferred in this study using LDhat. We drew the fitness effects of mutations from the common marmoset DFE inferred by Soni et al. (2025c). This DFE is comprised of four fixed classes (Johri et al. 2020): effectively neutral mutations (2*N*_ancestral_ *s* < 1, where *N_ancestral_* is the ancestral population size and *s* is the reduction in fitness of the mutant homozygote relative to wild-type), *f_0_* = 0.3; weakly deleterious mutations (1 ≤ 2*N*_ancestral_ *s* < 10), *f_1_* = 0.1; moderately deleterious mutations (10 ≤ 2*N*_ancestral_ *s* < 100), *f_2_* = 0.1; and strongly deleterious mutations (100 ≤ 2*N*_ancestral_ *s*), *f_3_* = 0.5. Chromosomal segments were then concatenated to create the entire non-PAR that could be scanned for selection. We performed genome scans for recent positive selection (using SweepFinder2 v.1.0; DeGiorgio et al. 2016) and balancing selection (using the *B_0MAF_* method implemented in BalLeRMix+_v1.py; Cheng and DeGiorgio 2020) on the simulated data. For SweepFinder2, we performed inference at each SNP using the command: *SweepFinder2 -lu GridFile FreqFile SpectFile OutFile*. For *B_0MAF_*, we performed inference at each SNP using the command: *BalLeRMix+_v1.py -I FreqFile --spect SfsFile -o OutFile --noSub –MAF –rec 1.456e-7*. We conservatively used the highest CLR value observed across all null model simulations as the null threshold for our empirical genome scans.

With these null thresholds in hand, we applied the above inference schema to our empirical X chromosome data. We curated candidate loci (i.e., those that exceeded null thresholds) by identifying genes under the likelihood surface and evaluating their functions and patterns of gene expression using the NCBI database (Sayers et al. 2022) and the EMBL-EBI Expression Atlas (Madeira et al. 2022).

## ACKNOWLEDGEMENTS

We would like to thank Eric Vallender for providing pedigree information for the individuals included in this study. Computations were performed on the Open Science Grid (OSG 2015) supported by the National Science Foundation Awards 2030508 and 2323298, and the Sol supercomputer at Arizona State University (Jennewein et al. 2023).

