## Supplementary Materials for "Inferring the relative contributions of evolutionary processes shaping X chromosome dynamics in the common marmoset (*Callithrix jacchus*) in the presence of twinning and hematopoietic chimerism"

| window size | min accessible length (bp) | no. of windows that meet threshold | proportion of total windows | 1.5 year generation time | | | 2 year generation time | | |
|---|---|---|---|---|---|---|---|---|---|
| | | | | 0.59 mya | 0.82 mya | 1.09 mya | 0.59 mya | 0.82 mya | 1.09 mya |
| 1Mb | 10,000 | 75 | 0.394737 | 2.92E-09 | 2.10E-09 | 1.58E-09 | 3.89E-09 | 2.80E-09 | 2.11E-09 |
| | 25,000 | 20 | 0.105263 | 3.04E-09 | 2.19E-09 | 1.64E-09 | 4.05E-09 | 2.92E-09 | 2.19E-09 |
| | 50,000 | 4 | 0.021053 | 2.75E-09 | 1.98E-09 | 1.49E-09 | 3.66E-09 | 2.64E-09 | 1.98E-09 |
| | 75,000 | 0 | 0 | - | - | - | - | - | - |
| | 100,000 | 0 | 0 | - | - | - | - | - | - |
| 100kb | 1,000 | 394 | 0.881432 | 2.39E-09 | 1.72E-09 | 1.29E-09 | 3.19E-09 | 2.29E-09 | 1.72E-09 |
| | 2,500 | 316 | 0.706935 | 2.61E-09 | 1.88E-09 | 1.41E-09 | 3.48E-09 | 2.51E-09 | 1.88E-09 |
| | 5,000 | 195 | 0.436242 | 2.82E-09 | 2.03E-09 | 1.53E-09 | 3.77E-09 | 2.71E-09 | 2.04E-09 |
| | 7,500 | 75 | 0.167785 | 2.59E-09 | 1.86E-09 | 1.40E-09 | 3.45E-09 | 2.48E-09 | 1.87E-09 |
| | 10,000 | 44 | 0.098434 | 2.54E-09 | 1.83E-09 | 1.38E-09 | 3.39E-09 | 2.44E-09 | 1.84E-09 |
| 10kb | 100 | 879 | 0.939103 | 2.35E-09 | 1.69E-09 | 1.27E-09 | 3.13E-09 | 2.25E-09 | 1.69E-09 |
| | 250 | 831 | 0.887821 | 2.30E-09 | 1.65E-09 | 1.24E-09 | 3.07E-09 | 2.21E-09 | 1.66E-09 |
| | 500 | 791 | 0.845085 | 2.35E-09 | 1.69E-09 | 1.27E-09 | 3.13E-09 | 2.25E-09 | 1.69E-09 |
| | 750 | 739 | 0.789530 | 2.33E-09 | 1.68E-09 | 1.26E-09 | 3.11E-09 | 2.24E-09 | 1.68E-09 |
| | 1,000 | 684 | 0.730769 | 2.37E-09 | 1.71E-09 | 1.28E-09 | 3.16E-09 | 2.27E-09 | 1.71E-09 |
| 1kb | 10 | 4,295 | 0.949381 | 1.98E-09 | 1.43E-09 | 1.07E-09 | 2.64E-09 | 1.90E-09 | 1.43E-09 |
| | 25 | 4,195 | 0.927277 | 1.86E-09 | 1.34E-09 | 1.00E-09 | 2.47E-09 | 1.78E-09 | 1.34E-09 |
| | 50 | 4,070 | 0.899646 | 1.73E-09 | 1.24E-09 | 9.35E-10 | 2.30E-09 | 1.66E-09 | 1.25E-09 |
| | 75 | 3,989 | 0.881742 | 1.72E-09 | 1.24E-09 | 9.30E-10 | 2.29E-09 | 1.65E-09 | 1.24E-09 |
| | 100 | 3,917 | 0.865827 | 1.71E-09 | 1.23E-09 | 9.27E-10 | 2.28E-09 | 1.64E-09 | 1.24E-09 |

**Supplementary Table S1.** Mean mutation rates on the common marmoset X chromosome for a number of window sizes, accessibility thresholds, generation times, and *C. jacchus*–*C. kuhlii* divergence times.

| male:female sex ratio | $\mu$ | $N_e$ based on simulated $\Theta_W$ | simulated $\Theta_W$ | $N_e$ based on empirical $\Theta_W$ |
|---|---|---|---|---|
| 0.3:0.7 | 2.16E-09 | 41,269 | 2.67E-04 | 58,767 |
| 0.4:0.6 | 2.22E-09 | 46,895 | 3.13E-04 | 56,963 |
| 0.45:0.55 | 2.26E-09 | 48,875 | 3.31E-04 | 56,102 |
| 0.5:0.5 | 2.29E-09 | 49,691 | 3.42E-04 | 55,266 |
| 0.55:0.45 | 2.33E-09 | 45,153 | 3.15E-04 | 54,455 |
| 0.6:0.4 | 2.36E-09 | 40,215 | 2.85E-04 | 53,668 |
| 0.7:0.3 | 2.43E-09 | 29,178 | 2.13E-04 | 52,159 |

**Supplementary Table S2. Sex-averaged per-site per-generation mutation rates, effective population sizes, and levels of genetic variation across simulated sex ratios.** The table shows the mean values of sex-averaged per-site per-generation mutation rates ($\mu$), effective population sizes ($N_e$), and levels of genetic variation ($\theta_w$) across 100 simulated replicates with different male:female sex ratios under the Soni et al. (2025b) demographic model, rescaled for the X chromosome (see "Materials and Methods" for details). $N_e$ was calculated using Watterson's $\theta_w$ (Watterson 1975) obtained from the simulations ($\theta_w = 3N_e\mu$) as well as from the empirical data ($\theta_w$= 3.8E-04).

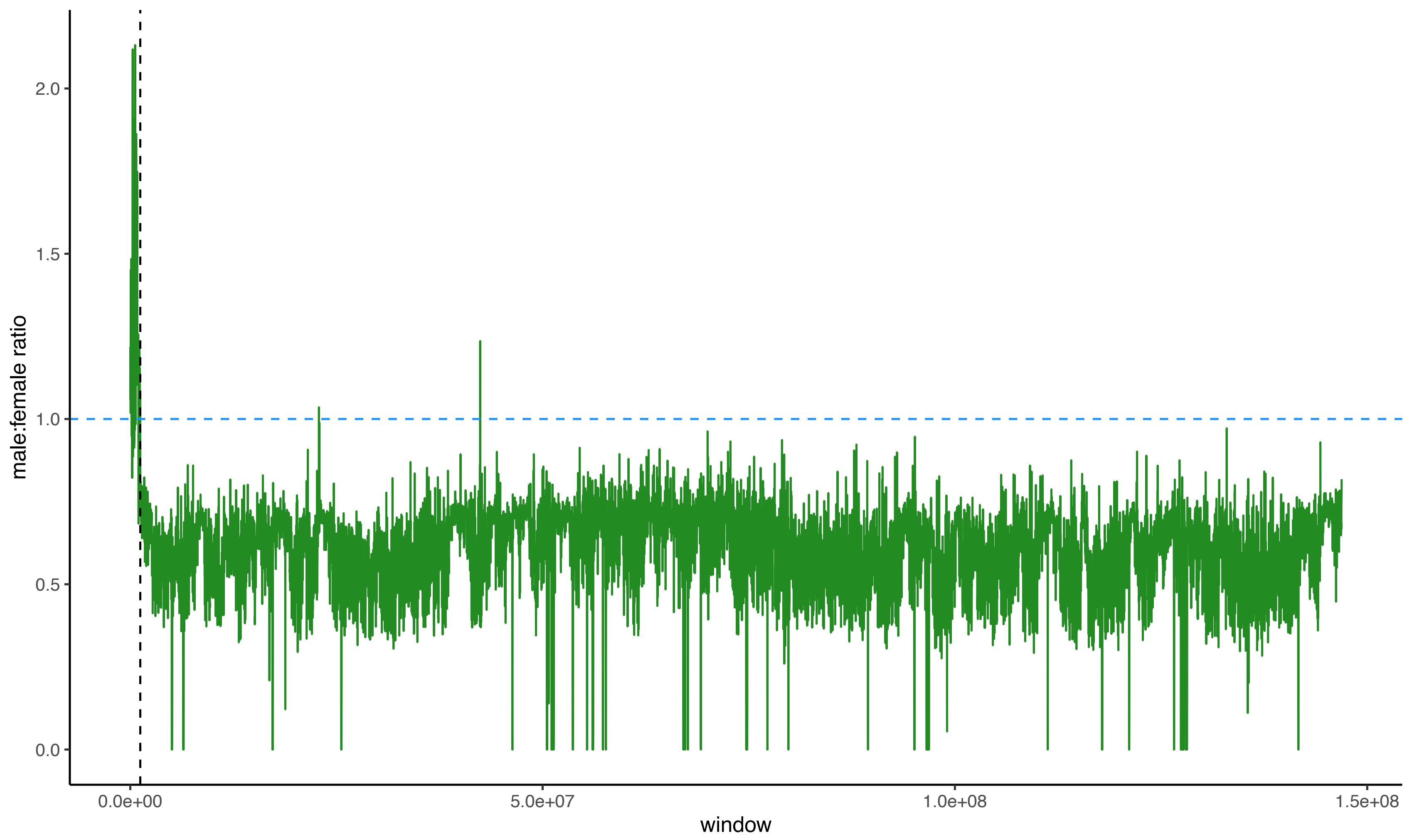


**Supplementary Figure S1:** Male:female ratios plotted across 10kb windows on the X chromosome in order to identify the pseudoautosomal boundary, with the horizontal blue dashed line indicating an equal ratio, and the vertical black dashed line indicating the identified pseudoautosomal boundary (and see Supplementary Figure S2 for a zoomed in version of the boundary region).

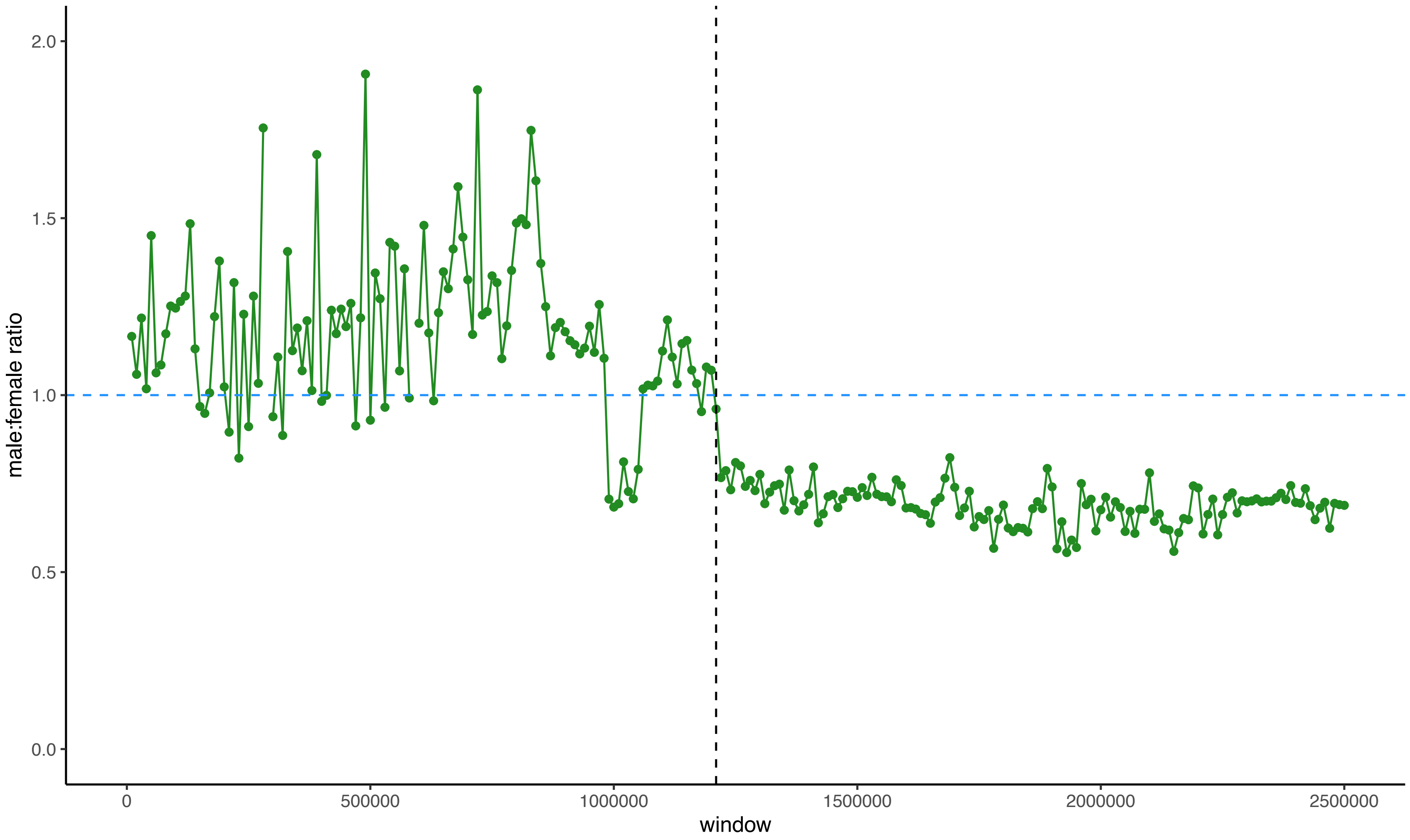


**Supplementary Figure S2:** Male:female ratios plotted across 10kb windows on the X chromosome in order to identify the pseudoautosomal boundary, with the horizontal blue dashed line denoting an equal ratio. Note that the x-axis is truncated at 2.5Mb (relative to Supplementary Figure S1) in order to better visualize the pseudoautosomal boundary (indicated by a vertical black dashed line).

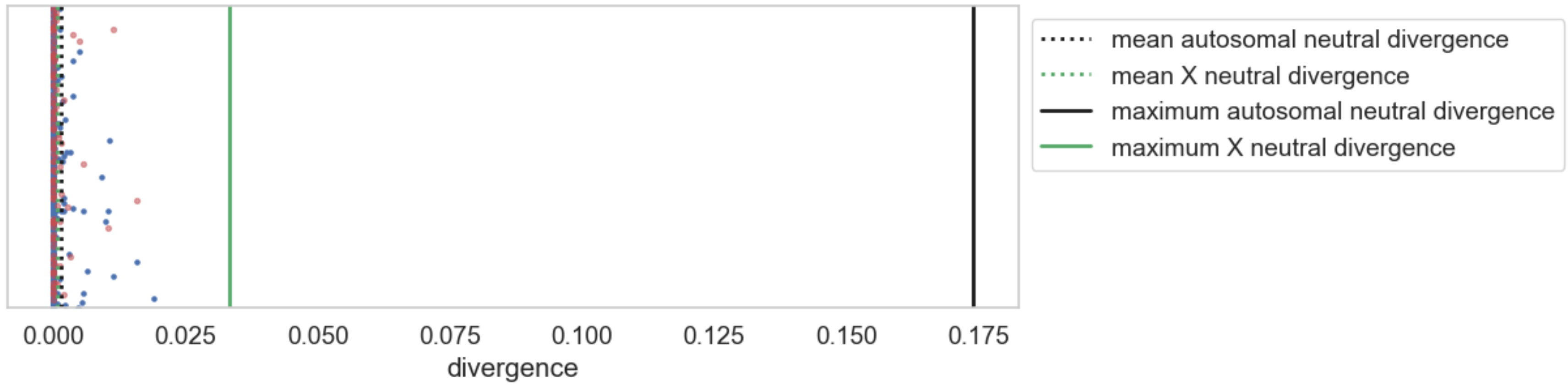


**Supplementary Figure S3:** Divergence across the X chromosome, with each point representing an X-linked exon. The mean and maximum neutral divergence on the X (green dotted and solid lines, respectively), and the mean and maximum autosomal neutral divergence (black dotted and solid lines, respectively) in 1kb windows are provided for comparison. The red dots are exons on the X, and the blue dots are exons on the autosomes.

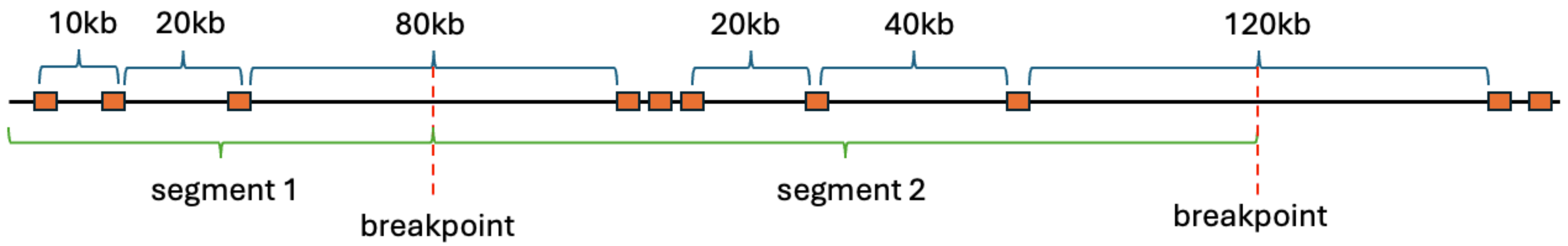


**Supplementary Figure S4. Schematic example of breakpoints for simulated segments used for generating null thresholds for genome scans.** A breakpoint is created wherever there is a >50kb distance between two exons, with the breakpoint occurring in the middle of this non-coding region.
